# The discriminator sequence is a primary determinant in the supercoiling response of bacterial promoters

**DOI:** 10.1101/2020.10.01.322149

**Authors:** Raphaël Forquet, Maïwenn Pineau, William Nasser, Sylvie Reverchon, Sam Meyer

**Affiliations:** Université de Lyon, INSA-Lyon, Université Claude Bernard Lyon 1, CNRS, UMR5240 MAP, F-69622, France

## Abstract

DNA supercoiling acts as a global transcriptional regulator, which contributes to the rapid transcriptional response of bacteria to many environmental changes. Although a large fraction of promoters from distant species respond to superhelical variations, the sequence or structural determinants of this behaviour remain elusive. Here, we propose the sequence of the “discriminator” element located downstream of the −10 hexamer to play an important role in this response, by modulating the facility of open-complex formation during transcription initiation. We develop a quantitative model of this regulatory mechanism relying on known parameters of DNA thermodynamics, and show that its predictions quantitatively match the *in vitro* and *in vivo* supercoiling response of stable RNA promoters previously measured, as well as the *in vivo* response of selected mRNA promoters with mutated discriminator sequences. We then test the universality of this mechanism by statistical analysis of promoter sequences in transcriptomes of phylogenetically distant bacteria under conditions of supercoiling variations, (1) by gyrase inhibitors, (2) by environmental stresses, (3) inherited in the longest-running evolution experiment. In all cases, we identify a robust and significant sequence signature in the discriminator region, suggesting that promoter opening underpins an ubiquitous regulatory mechanism in the prokaryotic kingdom, based on the fundamental mechanical properties of DNA.

## Introduction

Bacteria encounter rapid changes of environmental conditions (availability of nutrients, physical or chemical stresses) to which they respond by quick and global modifications of their transcriptional programme. Inspired by early studies, current mechanistic models of this regulation are mostly based on transcription factors (TFs) which bind at specific promoters and interact with RNA Polymerase (RNAP). Yet more than half of *E. coli* promoters are not targeted by any known TF [1], and entire organisms are even almost devoid of them [2, 3] but exhibit nonetheless a complex regulation. Global transcriptional control has been further explained by variations in RNAP composition (sigma factors, [4]) or abundance [5] depending on growth conditions, as well as RNAP-binding regulatory molecules such as ppGpp [6].

Besides this variability of the transcription machinery, the physical state of the DNA template itself is subject to cellular control through DNA supercoiling (SC), *i.e.*, the over- or under-winding of the double-helix by the action of topoisomerase enzymes and architectural proteins [7, 8, 9]. In bacteria, the chromosome is maintained at a negative SC level by the action of the DNA gyrase, which changes in response to environmental cues [9]. This level was soon discovered to affect the expression of many promoters, both *in vitro* and *in vivo* [10, 11, 12, 13, 14]. Mechanistic studies showed that, besides modulating the binding of regulatory proteins, it could influence the activity of RNAP itself, and could thus act as a global transcriptional factor [7, 8, 9]. Accordingly, whole-genome analyses of the transcriptional response to DNA relaxation induced by gyrase inhibitors exhibited a broad response, providing lists of “supercoiling-sensitive genes” [15, 16, 17, 3, 18].

In spite of its importance, no sequence or structural signature was ever clearly identified in support of the latter property. A possible reason is that SC affects transcription at many successive steps of the process, e.g., open-complex formation [19, 20], promoter escape [10], elongation and termination [21], and their combined action eluded the identification of simple determinants of supercoiling-sensitivity. Additionally, transcription in turn affects the local level of SC [22], and consequently, the response of a given promoter depends quite strongly on its genomic and physiological context [23, 24]. Altogether, the complexity of the interaction between SC and transcription explains why there still are no models able to predict, even qualitatively, the response of a given promoter to variations of SC [9].

One particular mechanism identified early as a putative strong factor in this response occurs at the step of open-complex formation during transcription initiation [25]. The unwinding of DNA strongly facilitates its denaturation, and thus the formation of the “transcription bubble” by RNAP [11]. Since this constraint affects all promoters, it may have a widespread effect on gene expression; yet the question then arises, how it may lead to transcriptional *regulation*, i.e., the *selective* activation/repression of a subset of promoters by *global* SC variations. An important observation was made when analysing several stable RNA promoters [26, 27, 19], which are both strongly SC-sensitive and subject to stringent control [28, 27, 19]. It was found that both properties are correlated to the presence of a G/C-rich “discriminator” sequence located between the −10 element and the transcription start site (TSS) [29] which is denatured in the open-complex. The discriminator has a variable length of 5-8 nucleotides (nt), does not harbour any consensus sequence but is bound by the *σ*1.2 domain of RNAP [30]. It was thus postulated that the unusually high G/C-content of these promoters affects the formation and stability of the open-complex, which may then be modulated by SC [7] or ppGpp [30]. But since most experimental studies were previously conducted on a few stable RNA promoters, it is not yet clear, if this regulation mechanism is a specificity of the latter, or a general regulatory principle also valid for the broader class of promoters of protein-encoding genes (mRNA promoters). It is the latter hypothesis that we consider in this paper.

In the following, (1) we develop a thermodynamic regulatory model of this principle, based on the free energy required to open the transcription bubble, and related to the G/C-content of the discriminator sequence. We show that it quantitatively reproduces the *in vitro* and *in vivo* SC-response of stable RNA promoters of *Escherichia coli* and *Salmonella enterica*; (2) we validate the same mechanism in selected mRNA promoters, by mutating their discriminator region and measuring their response to variations of SC *in vivo*; (3) we develop a statistical analysis of genome-wide expression data obtained after DNA relaxation by gyrase inhibitors, and show that the discriminator is the primary location of global promoter selectivity in these conditions; (4) we show that this sequence determinant is robustly detected in a series of distant bacterial species; (5) finally, we analyse this contribution in physiologically relevant conditions involving SC changes, induced either transiently in response to environmental stress, or inheritably in the longest-running evolution experiment. Altogether, this study highlights the discriminator as a global sensor of SC variations, and suggests that the thermodynamics of DNA denaturation is used as a basal and ubiquitous regulatory principle in the prokaryotic kingdom.

## Results

### Regulatory effect of the discriminator sequence in stable RNA promoters

Negative SC destabilises the double-helix and facilitates the melting of the transcription bubble during open-complex formation. This dependence is shown on Fig. 1B for the *tyrT* promoter of the tyrosine tRNA operon, one of the first SC-sensitive promoters described [26]. This curve is obtained from a physical model of DNA thermodynamics relying on knowledge-based enthalpic and entropic parameters of all base sequences [31, 32], using the precise 14-nt sequence of the region denatured in the open-complex (mostly encompassing the −10 hexamer and the discriminator, see Methods). Variations of the SC level should then directly affect the opening facility of promoters and thus their expression, and such a dependence was indeed observed for the *tyrT* promoter (blue) in both *in vitro* (Fig. 1C) or *in vivo* (Fig. 1D) transcription assays [20].

**Figure 1:**
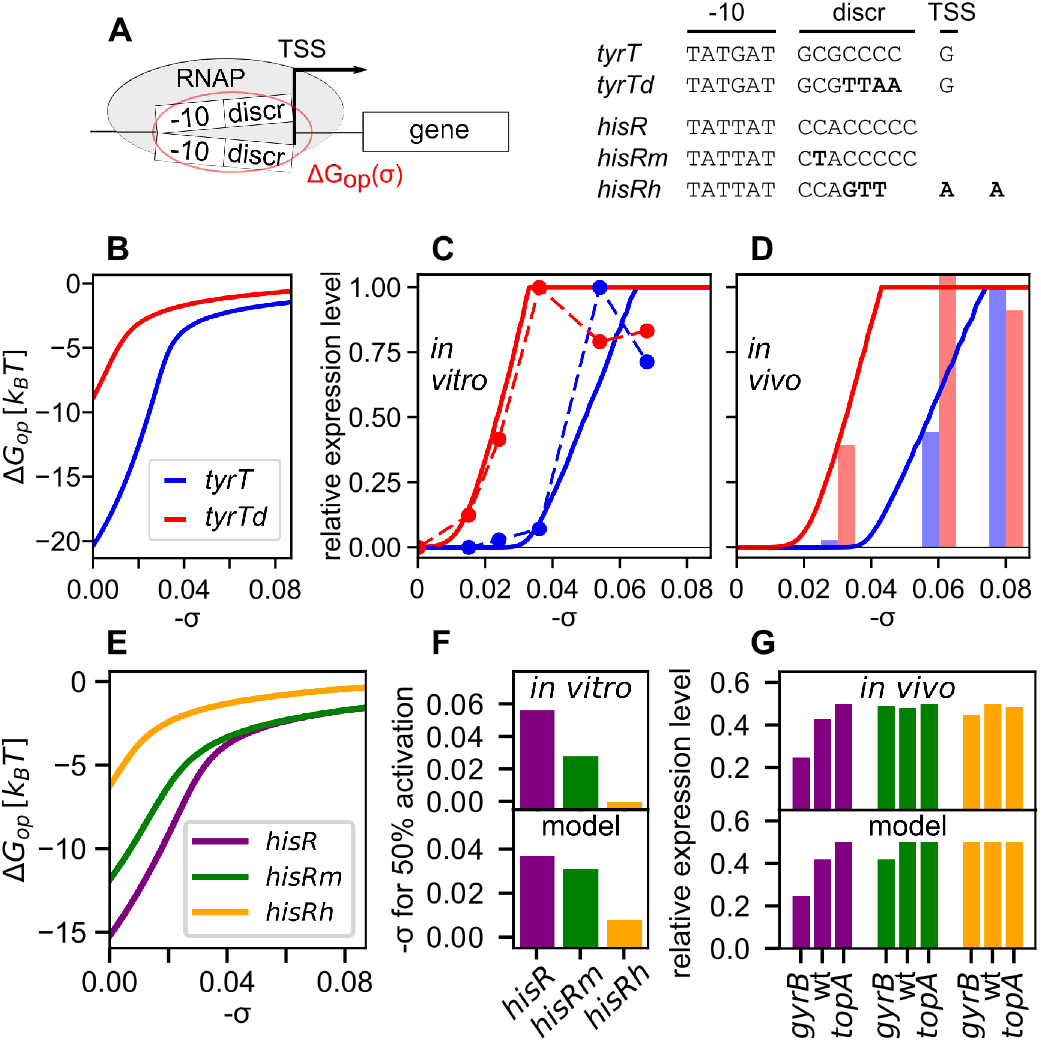
**(A)** Sequences from wild-type *tyrT* and *hisR* promoters, the mutant *tyrTd* promoter with A/T-rich discriminator [20], and the mutants *hisRm* and *hisRh*, with 1 and 5 substitutions C/G → A/T in the discriminator, respectively [19]. For *hisRh*, a shift in the transcription start site (TSS) (3 nt up-stream) was observed. **(B)** Transcription bubble opening free energies of *tyrT* and *tyrTd* promoters. **(C)** Transcription model predictions (solid lines) compared to the *in vitro* (dots) and **(D)** *in vivo* (bars) expression data from [20]. Data and computed values of the *tyrT* promoter are shown in blue, and those of *tyrTd* in red. **(E)** Transcription bubble opening free energies of *hisR*, *hisRm* and *hisRh* promoters. **(F)** Transcription model predictions compared to the *in vitro* and **(G)** *in vivo* expression data from [19]. Data and computed values of the *hisR*, *hisRm* and *hisRh* promoters are shown in purple, green and orange, respectively.

Further, the denaturation energy of the transcription bubble is known to be strongly dependent on the proportion of G/C bases. While the A/T-rich sequence of the −10 hexamer is relatively constrained due to its role in promoter recognition by the sigma factor, replacing four C/G by A/T nucleotides in the discriminator (*tyrTd* mutant) indeed strongly shifts the opening curve to the left (Fig. 1B, red curve), *i.e.*, favours DNA opening already at weaker SC levels. Strikingly, the resulting transcriptional activation curves (Figs. 1 C and D) closely follow the thermodynamic predictions.

This combination of quantitative expression data and computational modelling supports the notion that the open-complex thermodynamics is critical in the supercoiling-sensitivity of stable RNA promoters [27], with an important role of the discriminator sequence [26, 20]. We translated this qualitative notion into a quantitative thermodynamic model of transcription [33], based primarily on the promoter DNA opening curves (Fig. 1B). This model was kept voluntarily as simple as possible, since this mechanism is only one of the multiple steps by which SC affects transcription. It involves a single adjustable parameter (representing an opening assistance by RNAP), yet is able to quantitatively reproduce both *in vitro* and *in vivo* activation curves of these promoters (solid lines in Figs. 1 C and D). Theoretical foundations of the model are described in the Methods section, while the biological relevance and limitations are developed in the Discussion; in particular, the observed reduction of *in vitro* expression at very high SC levels (rightmost datapoints) likely reflects more complex mechanisms, such as the kinetics of promoter escape [10] or thermodynamic competition with structural transitions at neighbouring sites, which are known to play an important role in the genome [31].

We further tested the model using a similar dataset collected independently, based on the promoter of *hisR*, the histidine transfer RNA of *S. enterica* [19]. *In vitro* (Fig. 1F), the expression increases with negative SC, both in the WT or in mutant promoters of variable G/Crichness in the discriminator, closely following the DNA opening curves of the associated sequences (Fig. 1E), and are thus reproduced by the model without any parameter adjustment. *In vivo*, only the native promoter was affected (Fig. 1G) in topoisomerase mutant strains exhibiting a global SC shift either in the direction of DNA relaxation (*gyrB* mutant) or SC increase (*topA*). This feature was reproduced using the experimentally measured SC levels of these strains [34], suggesting that the two A/Tricher mutant promoters have reached a plateau where the denaturation energy, and hence the expression level, is almost independent of SC.

### Validation of model predictions on mutant mRNA promoters

While the role of the discriminator in SC-sensitivity has been previously described only in the case of stable RNA promoters [27, 19, 20], the physical constraints associated to the open-complex formation should equally affect mRNA promoters. We therefore tested if SC variations might be associated to a preferential activation/repression of mRNA promoters depending on the A/T-richness of their discriminator. Two families of synthetic promoters with mutated discriminators were constructed (Fig. 2A and Supplementary Tab. S1), based on (i) the *pheP* promoter of *E. coli*, which is SC-sensitive [16, 15] and is not regulated by any identified TF [1], and (ii) the *pelDpelE* genes of the enterobacterial phytopathogen *Dickeya dadantii*, which are paralogous virulence genes encoding similar pectinolytic enzymes, are directly regulated by more than ten TFs and are both supercoiling-sensitive [35] but harbour different discriminators. Promoters were fused on plasmids in front of a luciferase reporter gene (Fig. 2A), and their expression was analysed in *E. coli* cells in a microplate reader after treatment by the gyrase inhibitor novobiocin (see Methods). The employed plasmids are well established as reflecting the average SC level of the chromosome [36], in particular during DNA relaxation by novobiocin [37, 35, 38].

**Figure 2:**
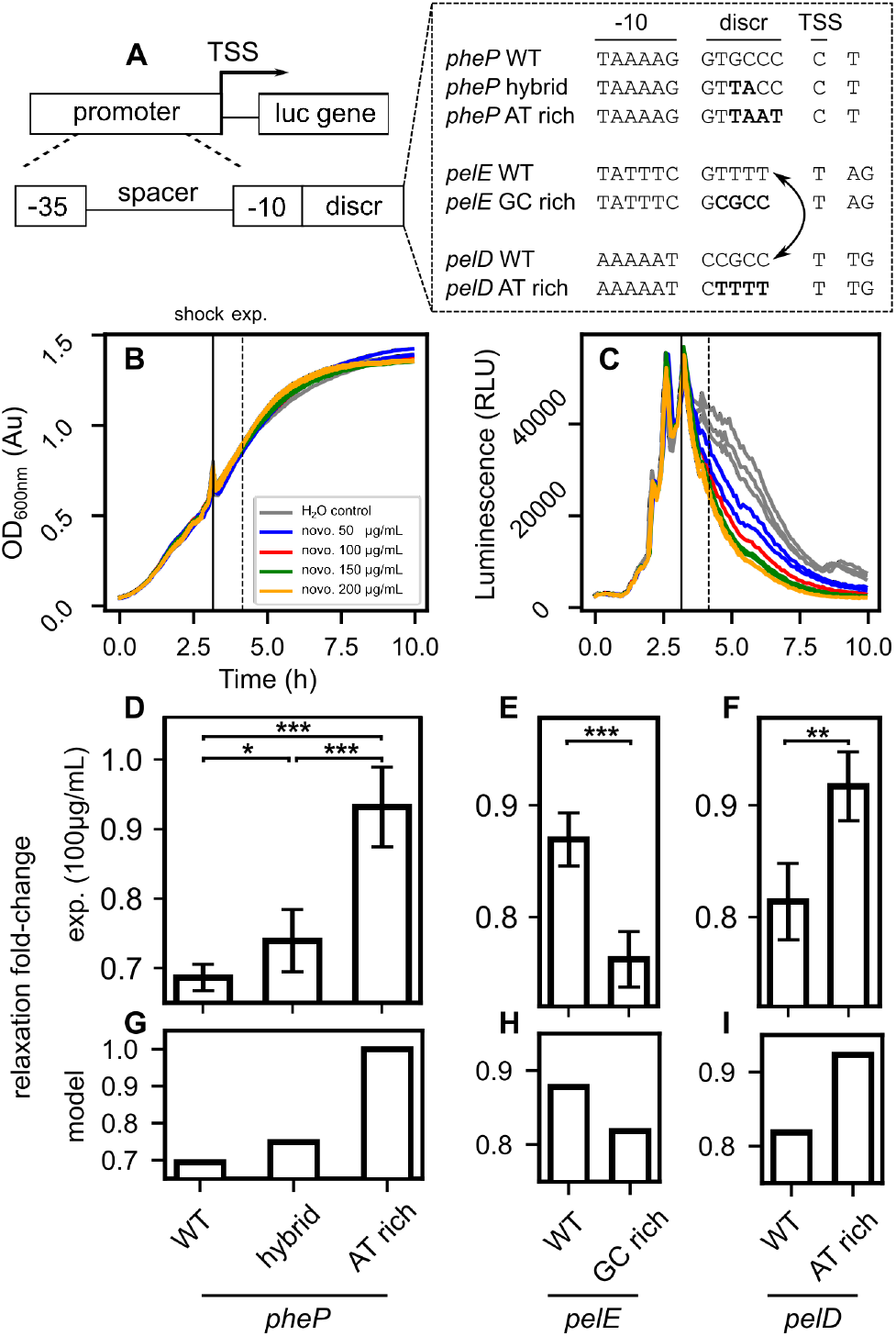
DNA relaxation response of mRNA promoters with mutated discriminators. **(A)** Promoter sequences were derived from *pheP* (*E. coli*) and *pelD*/*pelE* (*D. dadantii*), with mutated discriminators of various G/C content. **(B)** Bacterial growth monitored in a microplate reader (*E. coli* bacteria carrying plasmids with *pheP* hybrid promoter in rich medium). A novobiocin shock was applied in mid-exponential phase (different sublethal concentrations shown). The slight increase at shock time is an optical artefact due to the opening of the recorder. **(C)** Expression of the *pheP* hybrid promoter monitored by luminescence (see all raw datapoints in Supplementary Fig. S2). **(D)** Relaxation fold-changes computed 60’ after novobiocin shock (100 *μ*g/mL) in *pheP*-derived promoters. As expected, the repression factor reduces with increasing A/T%. **(E)** The DNA relaxation response of *pelE* and **(F)** *pelD* are reversed when a tetranucleotide is swapped between their discriminators, with low and high G/C content respectively. **(G)** Relaxation fold-changes predicted by the model reproduce the experimental observations on *pheP*-derived promoters, as well as **(H)** *pelE*-and **(I)** *pelD*-derived promoters, assuming a weak relaxation compatible with the observed repression levels (see Materials and Methods).

We first checked that the presence of the plasmids did not affect bacterial growth, and that the expression patterns of two promoters as well as their response to novobiocin were consistent when inserted either in plasmid-borne or in chromosomal luciferase fusions (Supplementary Fig. S1). These observations match previous similar comparisons involving other promoters and plasmids [39, 40], and confirm that the reduction in luminescence observed following the shock (raw data in Supplementary Fig. S2) is due to SC-dependent transcriptional regulation rather than plasmid-specific effects. We then compared the relative effect of the novobiocin shock on the different plasmid-borne promoters. For the *pheP*-derived promoters (Fig. 2D), cells were grown in LB medium. As expected, we found that the expression fold-change (treated vs non-treated wells) was strongest for the native G/C-rich promoter, significantly reduced for the hybrid promoters (with two mutated nucleotides in the discriminator), whereas the A/T-rich discriminator (with four mutated nucleotides) was almost insensitive to DNA relaxation. Similarly, swapping four nucleotides between the discriminators of *pelE* and *pelD* (Fig. 2 E and F) strikingly reversed their response to DNA relaxation in minimal medium. The relatively modest (but highly significant) repression levels are partly due to a buffering effect of the reporter system, and accordingly, all results are reproduced with the model assuming a weak global relaxation magnitude (Fig. 2 G-I and Methods), where the direction of the promoters’ predicted response is inscribed in their sequences and is therefore robust when this magnitude is varied.

These results show that the discriminator sequence is an important element in the supercoiling response, not only of stable RNA promoters, but also of mRNA promoters of diverse biological functions and regulation complexities, and this effect is detectable in different physiological conditions (rich vs minimal medium). Since the proposed mechanism of open-complex formation is involved in RNAP-promoter interaction independently from additional regulatory proteins, we now enlarge the scale of the analysis to entire genomes.

### The discriminator is the primary location of promoter selectivity by DNA relaxation

We first looked at the variability of discriminator G/C-contents among mRNA promoters in various species, based on available TSS maps (Supplementary Fig. S3). These distributions are wide, and alike *pheP* and *pelE*, a large class of promoters have G/C-rich discriminators comparable to those of stable RNAs. Based on the previous analysis, we hypothesised that such promoters would be more repressed by a DNA relaxation induced by gyrase inhibitors than those harbouring an A/T-rich discriminator. However, in contrast to the mutation studies above, here the compared promoters differ by many additional factors beyond their discriminator sequence (upstream and downstream sequences, genomic context, NAP binding, transcriptional regulators, etc) which may contribute to their supercoiling response, and we therefore looked for a statistical relation rather than a prediction valid for all analysed promoters.

We aligned all *σ*70 promoters of *Salmonella enterica*, and looked at their average A/T% profile (Fig. 3A) depending on their response 20 minutes after a novobiocin shock [18]. Strikingly, although this content exhibits a characteristic non-uniform pattern along the promoter (with an expected peak at the −10 element), the signals of the two groups of promoters are indistinguishable everywhere except in the region between −10 and +1, precisely where we expected the observed difference (*P* < 10^−5^ around position −2, Supplementary Tab. S2). This observation, obtained independently from the mutation studies above, confirms that the discriminator region is the primary location of selectivity for the relaxation response. As a comparison, no significant difference is detected at the −10 element, suggesting that this selectivity is not related to a difference in sigma factor usage. Further, classifying the promoters based on their discriminator sequence composition (Fig. 3B) exhibits a clear and highly significant (approximately linear) effect on the proportion of activated promoters (correlation *P* - value < 10^−4^).

**Figure 3:**
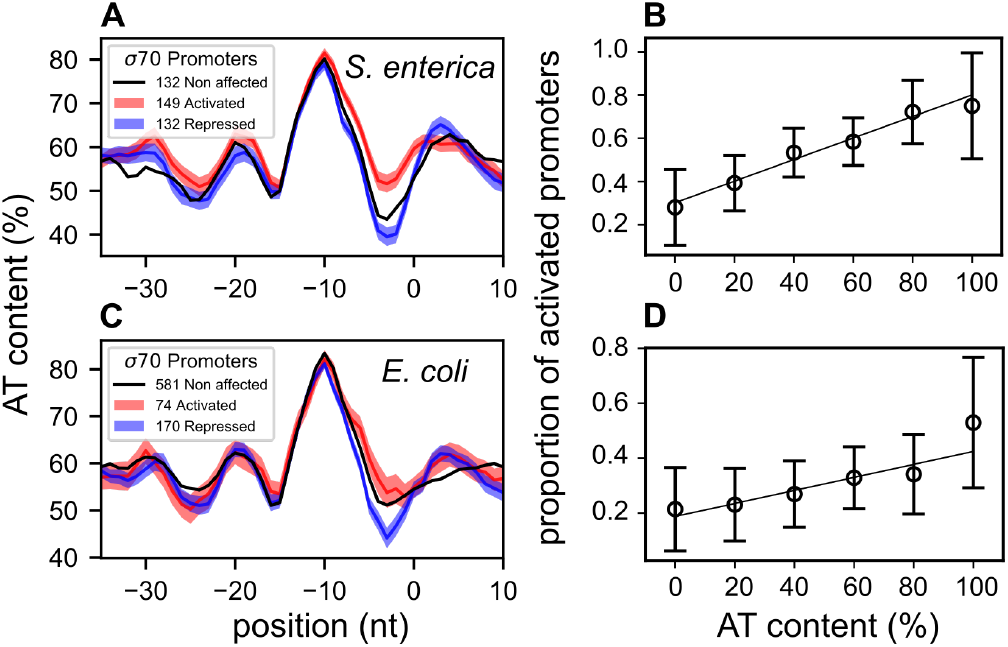
Genome-wide relation between discriminator sequence and promoter selectivity during DNA relaxation. **(A)** Average A/T% profiles of *S. enterica σ*70 promoters along 5-nt-centred windows, depending on their response to novobiocin-induced chromosomal relaxation [18]. The profiles are very similar (shaded areas represent one standard error) except in the discriminator region (between −10 and +1 positions). **(B)** Proportion of activated promoters among those responsive to DNA relaxation, depending on their A/T% in a 5-nt window around position −2. The resulting linear regression is highly significant (*P* < 10^−4^). **(C)** Average A/T% profiles of *E. coli σ*70 promoters depending on their response to norfloxacin-induced DNA relaxation (LZ54 vs LZ41 strains, [16]). The resulting pattern is very similar to that observed in *S. enterica* in spite of strong differences in protocols. **(D)** Same as B, for the *E. coli* data (*P* = 0.011).

### A robust relation observed across distant bacterial phyla

Since the investigated mechanism relies on highly conserved molecular actors, RNAP and topoisomerases, it might affect a particularly broad range of bacterial species. We therefore tested if our observations can be reproduced in other organisms. Transcriptomic data were obtained in *E. coli* with DNA microarrays after norfloxacin shock in two alternate topoisomerase mutant strains [41], resulting in a strong magnitude of DNA relaxation [16]. In spite of strong differences in the experimental protocol compared to the *S. enterica* dataset, the obtained pattern is remarkably similar (Fig. 3C and D). Importantly, whereas in the first experiment (treated vs non-treated cells), this pattern might include contributions from SC-independent drug-response pathways, here the two compared samples received exactly the same treatment, and any such unwanted contribution should thus not be apparent. The slightly weaker observed effect might also be due to the lower sensitivity of the employed transcriptomic technology.

In the enterobacterium *D. dadantii*, the response to relaxation by novobiocin was monitored in minimal medium [24], based on identified gene promoters [42], exhibiting the same pattern (Fig. 4C) as in *E. coli* (Fig. 4A) and *S. enterica* (Fig. 4B). At a drastically larger evolutionary distance, in the cyanobacterium *Synechococcus elongatus*, SC was shown to be a major determinant of the circadian oscillatory genomic expression [43]. The transcriptomic response to DNA relaxation was not monitored directly, but the phasing of gene expression in this oscillation can be used as an indirect proxy of this response [43], although many other metabolic signals may be equally correlated and could contribute to this signal. The promoter analysis yields a similar difference of discriminator sequence (Fig. 4D), of slightly lower magnitude possibly due to these additional regulatory mechanisms and a poorer definition of promoters (and associated sigma-factors). Finally, the response to novobiocin was also monitored in the small tenericute *Mycoplasma pneumoniae* [3], in which transcriptional regulation is poorly understood due to the quasi-absence of TFs [44]. Although the signal is here possibly also weakened by the lower number of promoters, it is still significant (Fig. 4E). Altogether, although these two latter species differ widely from the others in terms of phylogeny, lifestyle, and average G/C content (in particular, *M. pneumoniae* has very few promoters with strongly G/C-rich discriminators, Supplementary Fig. S3), these robust results consistently suggest that the ancestral infrastructural constraint of DNA opening, coupled with the conserved activity of topoisomerases, indeed underpins a global regulatory mechanism throughout the prokaryotic kingdom.

**Figure 4:**
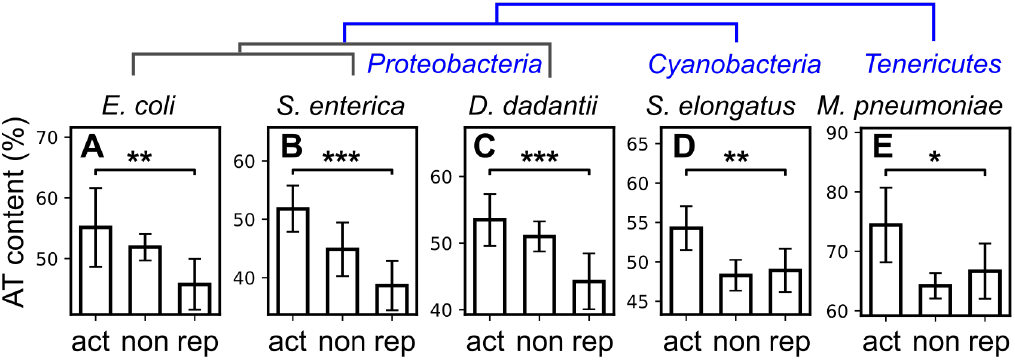
A robust statistical relation between discriminator A/T% and promoter’s response to DNA relaxation by novo-biocin (act: activated, non: no significant variation, rep: repressed) is observed in distant bacterial species: **(A)** *E. coli* (*P* = 0.010, relaxation by norfloxacin in LZ54 vs LZ41 mutant strains) [16]; **(B)** *S. typhimurium* (*P* < 10^−5^); **(C)** *D. dadantii* (*P* < 10^−3^); **(D)** *S. elongatus* (*P* = 0.004); **(E)** *M. pneumoniae* (*P* = 0.029). In enterobacteria, only *σ*70 promoters were considered, and aligned at the −10 element. In the two other species, all promoters were aligned at their annotated TSS. A/T% are computed in a 5-nt window in the discriminator (see Materials and Methods). A schematic phylogeny is depicted above.

### Global response to stress conditions and inheritable supercoiling variations

While sublethal antibiotic shocks are the classical method of choice to specifically induce rapid DNA relaxation [47, 9], in natural conditions the latter is rather triggered by sudden changes of environmental conditions, especially by physico-chemical stress factors like temperature, acidity, oxidative agents etc. The resulting rapid SC variations were found to be conserved even in distant species, e.g., increase of negative SC by cold shock, DNA relaxation by heat shock or oxidative stress [9]. We therefore tested if the sequence signature expected from the analysis above might be detected in published transcriptomic data.

Temperature shocks provide a useful example, since heat and cold shocks both put the bacteria under stress, while affecting the SC level in opposite directions (relaxation [48] and overtwisting [49] respectively, Supplementary Tab. S2). The analysis of the corresponding transcriptomic datasets obtained in independent studies clearly confirms the expectations, with G/C-rich discriminators being respectively repressed and activated with a linear dependence in the sequence content (Figs. 5 A to D). In the case of oxidative stress associated to DNA relaxation (induced by H_2_O_2_), the response was analysed in the enterobacteria *E. coli* and *D. dadantii* [50, 35], where the pattern is indeed very similar and matches the expectations. Altogether, this analysis suggests that, beyond stress-specific regulation pathways mediated by dedicated regulatory proteins, the SC variations induced in these conditions play a direct role in the resulting global reprogramming of gene expression by modulating the RNAP-promoter interaction through the discriminator sequence.

**Figure 5:**
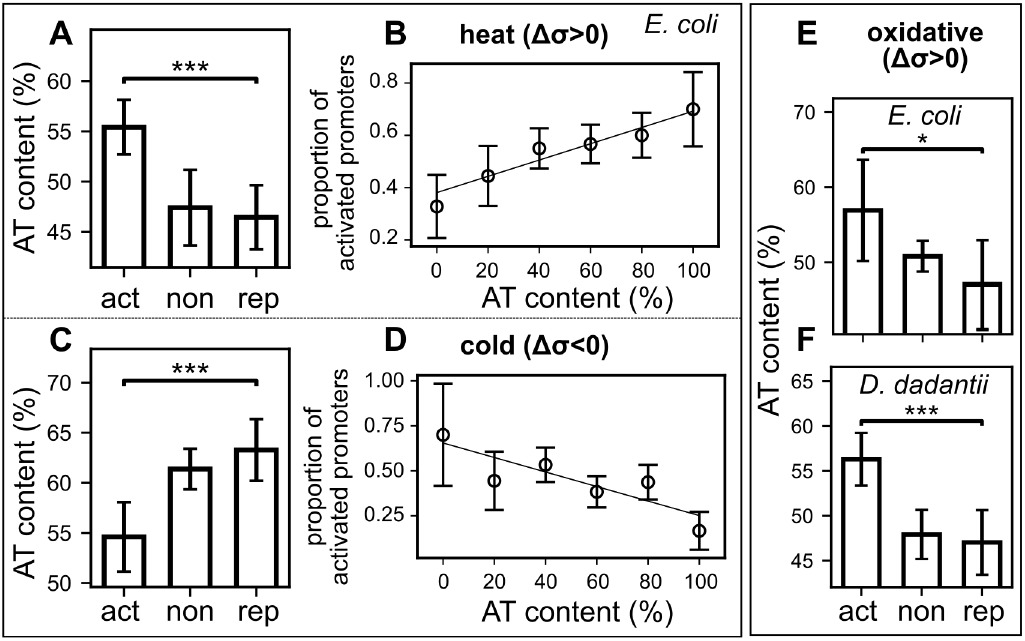
Relation between discriminator sequence and response to SC variations induced by environmental stress conditions. **(A)** During heat shock in *E. coli* [45] triggering a transient DNA relaxation (Δ*σ*>0), activated promoters have discriminators with higher A/T% than repressed ones (*P* < 10 – 5) as expected from the presented model. **(B)** The proportion of promoters activated by heat shock increases linearly with A/T content (correlation *P* = 0.007, same data as A). **(C)** In a cold shock in *E. coli* [46] inducing an opposite SC variation (increase in negative SC, Δ*σ*<0), the relation is reversed, with a preference of G/C-rich discriminators among activated promoters (*P* < 10^−4^), as we expected. **(D)** The promoters’ response is linear and significant (correlation *P* = 0.026, same data as C). **(E)** Same as A, during an oxidative shock in *E. coli* [46] inducing DNA relaxation (*σ*>0, *P* = 0.017). **(F)** Same in *D. dadantii* (*P* < 10–4) [17] where the shock was shown to induce the same SC response [35], showing the conservation of the mechanism.

Finally, we address the question, if the investigated mechanism might be involved not only in transient responses, but also in inheritable modifications of the expression programme. In the longest-running evolution experiment with *E. coli* [51], point mutations inducing variations of the SC level were indeed quickly and naturally selected [52], as they provided substantial fitness gains that were attributed to the resulting global change of the transcriptional landscape [24]. In the investigated conditions of growth in nutrient-poor medium, a first mutation (in *topA*, among 6 in total) before 2000 generations, and a second mutation (in *fis*, among 45 in total) before 20,000 generations both lead to an inheritable increase of negative SC (Fig. 6A). Based on the modelling, these mutations should predominantly enhance the expression of promoters with G/C-rich discriminators in the evolved strains. Such a tendency is indeed observed, both after 2000 generations where the signal is strongest (Fig. 6B, *P* = 0.005) and after 20,000 generations (Fig. 6C, *P* = 0.011, Supplementary Tab. S2), where 43 accumulated mutations besides these two affecting SC probably contribute in rewiring the regulatory network and blurring the signal. The detected signal suggests that the proposed biophysical regulatory mechanism is not only involved in rapid changes of gene expression, but may be used as a driving force in the evolution of genomes.

**Figure 6:**
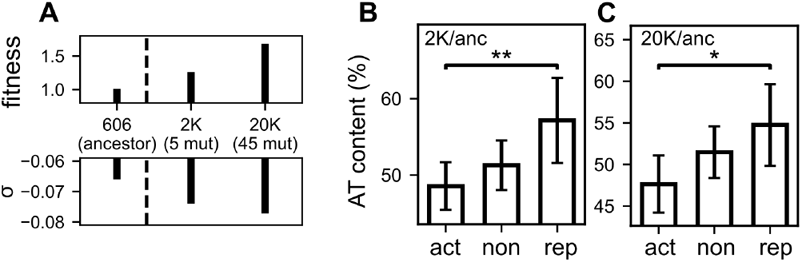
**(A)** In the longest-running evolution experiment [51], two point mutations naturally acquired by *E. coli* [52] induced successive increases of negative SC: one in *topA* before 2000 generations (among the five observed), and one in *fis* before 20000 generations (among the 45 observed), and are associated to fitness gains through modifications of global gene expression [24]. **(B)** Proportion of A/T content in the discriminator of promoters activated, repressed or unaffected in the evolved strain 2K compared to the ancestor. As expected from our modelling for an increase in negative SC, activated promoters with G/C-rich discriminators are more activated (*P* = 0.005). **(C)** In the 20K evolved strain, the same difference is observed (*P* = 0.011) although less significant, possibly due to many other mutations affecting the regulatory network.

## Discussion

While the influence of the discriminator sequence in the supercoiling sensitivity of stable RNA promoters has been reported for decades [26, 27, 19, 20], no sequence determinant was ever clearly identified in the broader class of supercoiling-sensitive mRNA promoters identified in various bacteria. In this work, we proposed a simple thermodynamic model of open-complex formation that quantitatively accounts for this regulation mode, and we showed that it applies not only to stable RNA promoters previously analysed, but also to mRNA promoters. A statistical analysis carried in various species and experimental conditions consistently and robustly showed that the discriminator sequence is a significant determinant of promoter supercoiling-sensitivity, and highlights the widespread relevance of this mechanism in the genome-wide response to transient or inheritable variations of SC levels.

### Quantification and limitations of the regulatory mechanism

A major difficulty when analysing SC-induced regulation is that it affects the transcription process at multiple steps, from the binding of regulators to the activity of RNAP itself during transcription initiation [10], elongation and termination [21]. While we focused our analysis on the discriminator sequence, the reader should keep in mind that many other mechanisms contribute in enhancing the complexity of this regulation: (1) the influence of DNA conformation on its interaction with regulatory proteins [9]; (2) competing structural transitions (denaturation, cruciform exclusion, G-quadruplex, Z-DNA) occurring in nearby regions depending on the SC level, and strongly affecting the SC-response at the initiation site [31]; (3) the modulation of the effective SC level available for denaturation, because of twist/writhe dynamics and local mechanical constraints imposed by regulatory proteins [9]; (4) the heterogeneity of SC levels in different topological domains along the chromosome [53]. These factors and others explain why, in the analysed transcriptomic data, the effect of the discriminator sequence emerges as a statistical feature at the genomic scale, rather than a predictive signal dictating the response of each individual promoter as observed in mutation studies. In particular, since a negative SC level favours the denaturation of G/C-rich as well as A/T-rich sequences (Fig. 1), this mechanism alone is insufficient to explain the existence of a class of relaxation-activated promoters, such as *gyrA/B* [13]. This behaviour might be explained by more complex mechanisms involving the kinetics of promoter opening and escape by RNAP, where the stability of the open-complex becomes unfavourable if it leads to abortive rather than processive transcription [10, 25], or by thermodynamic competition with other structural transitions occurring at nearby sites [31].

In spite of these limitations of our simple modelling, and based on the sequence signal observed in transcriptomic data, can we quantify the contribution of this specific mechanism in the genome-wide supercoiling response? To estimate this magnitude, we developed a genome-wide prediction of the relaxation-response based solely on the thermodynamic opening model developed above (independently from all other transcriptional effects of SC), and computed the proportion of accurate predictions among the observed differentially expressed genes (activation or repression). Compared to a null (random) model, this proportion is improved by around 10-15% of the responsive genes in the investigated relaxation and environmental stress assays (usually several hundreds, representing a high statistical significance of predictive power, see details in Supplementary Tab. S2). Considering the many alternate regulatory mechanisms by SC, for which no comparable estimates are available at the genomic scale (most of them lacking quantitative models), this proportion computed from a single step without any parameter adjustment is quite notable. Additionally, it is likely underestimated because of many inaccurately annotated promoters (a single-nucleotide resolution is required but not always achieved). Note that, because the total mRNA levels is normalised in transcriptomic data (predefined sequencing depth, erasing any global activation/repression effect), we introduced a comparable normalisation step in the computation: as a result, a fraction of A/T-rich promoters appear as activated by the DNA relaxation even if they are more difficult to open by RNAP (by competition with G/C-rich ones, see Methods).

### Simultaneous regulation by SC and ppGpp at the discriminator

Among various further regulatory mechanisms related to this study, the alarmone ppGpp, classically associated to the stringent (starvation) response [6], deserves special attention. In contrast to many TFs, ppGpp affects the expression of a large subset of the genome by binding RNAP in combination with the transcription factor DksA [54], and modulating the stability of the open-complex [27]. Its repressive effect is not dependent on a strict sequence motif, but rather on the presence of a C nucleotide at position −1 [54]. This regulatory mechanisms thus presents many similarities with the one investigated here, and both are involved in the regulation of bacterial growth, thus raising the possibility of an interplay between these two pathways [55, 27]

We first checked that the sequence signatures identified in this study were not due to a regulatory effect involving ppGpp rather than SC. It was observed that gyrase inhibition does not trigger any growth arrest (Fig. 2B) nor signature of stringent response [16], and accordingly, an analysis of the expression levels of genes involved in ppGpp synthesis (*gppA*, *spoT*, *relA*) does not exhibit any significant response [15, 16, 17, 18, 3]. Thus, DNA relaxation does not trigger ppGpp production, and even if the two pathways are associated to a similar sequence signal in the discriminator, the observations made in this study are indeed due to a ppGpp-independent effect of SC.

We then carried a sequence analysis of the promoters directly regulated by ppGpp through its binding to RNAP, as identified at the genomic scale in a recent study in *E. coli* [54]. As expected, a strong difference in G/C% between the many promoters activated and repressed by ppGpp induction (representing 70% of *σ*70 promoters in total) is detected in the discriminator (Supplementary Fig. S4A), similar to the pattern observed with DNA relaxation (Fig. 3), confirming that the two pathways affect transcription at the whole-genome scale based on similar promoter sequence determinants.

While DNA relaxation does not induce ppGpp production, it was conversely shown that the induction of high levels of ppGpp by the stringent response does trigger a sharp fall in SC levels in *E. coli* [27]. It is thus plausible that the strong sequence signature observed after ppGpp induction (Supplementary Fig. S4A) actually results from the addition of two independent factors of open-complex destabilisation: RNAP binding by ppGpp, and DNA relaxation. Interestingly, the transcriptional response to ppGpp induction was also monitored in mutant cells where it is unable to bind RNAP, thus inhibiting its direct regulatory activity [54]. Remarkably, almost half as many genes respond as in the wild-type cells (representing 35% of *σ*70 promoters, although with weaker magnitudes and slightly slower response times), and these promoters exhibit a similar (albeit weaker) sequence signature at the same location (Supplementary Fig. S4B). A plausible explanation is that ppGpp induction indeed triggered a DNA relaxation [27], resulting in a similar but partial response compared to wild-type cells. This scenario remains hypothetical as the SC levels were not directly measured in these samples; it would then likely involve a post-transcriptional effect of ppGpp on gyrase activity, as frequently occurs in response to stress or metabolic signals [9].

Altogether, this combined analysis of transcriptomic data fully confirms the notion that the regulation by SC relaxation and ppGpp are distinct but partially redundant in their transcriptional effect. More precisely, SC relaxation may be considered as a more fundamental form of regulation relying on the basic infrastructure of transcription, whereas ppGpp synthesis may itself trigger DNA relaxation (but not conversely). The relationship between the two pathways is further emphasised by the observation that, in the evolution experiment, the two genes most quickly and robustly affected by mutations are *topA* and *spoT* [56, 52], involved precisely in SC and ppGpp synthesis/degradation [6] respectively. Interestingly, the *spoT* mutation alone explains only a part of the observed transcriptional change [57], while similarly, the *topA* mutation alone generates only a fraction of the observed signal at the discriminator (Supplementary Fig. S5), suggesting a synergistic action of these two mutations [56, 52]. The additive selection of promoters based on the same sequence signal at the discriminator provides a plausible and natural mechanistic explanation for this feature.

Finally, in the datasets obtained with environmental stress conditions that we have analysed (Fig. 5), the genes associated to ppGpp synthesis are partly responsive, but rather in an opposite direction to the discriminator sequence signature observed (repression in heat and oxidative stress, slight activation in cold stress) and this pathway does probably not contribute significantly to the observed signal.

## Methods

### Synthetic promoters

230, 329 and 313 nt sequences upstream of the *pheP*, *pelE* and *pelD* start codons, respectively, were synthesised with mutations in the discriminator (GeneCust) and individually cloned into pUCTer-*luc* plasmids (Supplementary Tab. S1) upstream of a luciferase reporter gene (*luc*). *E. coli* strain MG1655 cells were then transformed with these plasmids using a standard electroporation procedure.

### Measurement of DNA relaxation response of mutant promoters *in vivo*

*E. coli* cells carrying the plasmids with the different promoters were recovered from glycerol stock (−80°C) and grown overnight (about 16h) on LB Agar plates at 37°C. The obtained colonies were further transferred to liquid cultures overnight (about 16h), with shaking at 200 rpm under selective antibiotic pressure (Ampicillin 60 μg/ml final). LB medium was used for bacteria carrying plasmids with *pheP*-derived promoters, whereas M63 minimal medium supplemented with 0.2% glucose was used forbbacteria carrying plasmids with *pelE* and *pelD*-derived promoters. Cells were washed (2’ centrifugation at 8000 rpm), then outgrowth cultures were performed in the same medium without antibiotics, stopped during exponential phase, and diluted for a final *OD*_600*nm*_ = 0.1 in a 96-wells microplate. Each well (200 μl final volume) contained the chosen medium supplemented with D-luciferin (450 μg/ml final). The microplate was placed in a humidity cassette and grown at 37°C until stationary phase was reached in a microplate reader (Tecan Spark). The *OD*_600*nm*_ and luminescence were measured every 5min, preceded by a 45s shaking step (double orbital, 3.5mm amplitude). During mid-exponential phase for *pheP* and early-exponential phase for *pelE* and *pelD*, the microplate was taken out and DNA relaxation was transiently induced by injecting 5 μL of novobiocin (50, 100, 150 and 200 μg/mL final concentrations tested) using a multichannel pipette. Data files produced by the microplate reader were parsed using a Python home-made script and the response to DNA relaxation was computed by comparing the luminescence values (in triplicates) of the novobiocin-shocked strain compared to the same strain injected with water (novobiocin solvent) 60’ after shock. The employed firefly luciferase has a short life-time, between 6’ in *B. subtilis* [58] and 45’ in *E. coli* [59]. Confidence intervals and p-values were computed using Student’s distributions.

### Genome-wide analyses of discriminator sequences

Transcriptomes obtained after DNA relaxation by antibiotics, inheritable supercoiling variations or environmental stresses were collected from the literature (Supplementary Tab. S2), as well as genome-wide TSS maps, and a scan for promoter motifs upstream of TSSs was conducted with bTSSfinder [60]. F or enterobacteria, *σ*70-dependent promoters were retained and aligned at their −10 site to reduce statistical noise. Some of them also bind other *σ* factors, but *σ*70 is predominant in exponential phase where the analysed samples were collected. The A/T% content was computed along 5-bp sliding windows (Fig. 3A and C). Promoters were classified according to their A/T% in a 5-nt window centred around position −2 rather than the entire discriminator (of variable size), which improves the statistical analysis while not affecting the distribution of promoters significantly (Supplementary Fig. S3). This relation was quantified either by linear regression (Fig. 3 and 5) or by a chi2 test between activated and repressed promoters (Fig. 4 to 6, Supplementary Tab. S2). For *Synechococcus elongatus* and *Mycoplasma pneumoniae*, all promoters were retained and aligned at their TSS; the resulting A/T% signal had a poorer signal definition and was shifted of around 5 nt (peak at the −10 element), and the same shift was applied to the position of the windows used in the analysis. All error bars shown are 95% confidence i ntervals, except the coloured shaded areas of Fig. 3 (67% confidence intervals).

### Model of transcriptional regulation by SC

The observed correlation between promoter opening thermodynamics and expression strength (Fig. 1) is accounted for by a thermodynamic regulatory model [33] involving a single adjustable parameter:

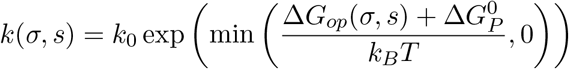

where *k* is the transcription rate, *k*_0_ the basal (maximal) rate, *s* is the precise 14-nt sequence of the denatured region in the open-complex [61] of estimated negative opening penalty Δ*G*_*op*_ (Fig. 1B), and *k*_*B*_*T* is the Boltzmann factor. 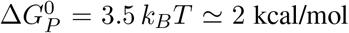 kcal/mol is a global energy parameter, representing an opening assistance by RNAP during open-complex formation, adjusted on the data of Fig. 1C (and kept constant for all promoters and species in the following computations). At high negative SC levels, the opening penalty becomes negligible (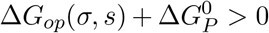, Fig. 1B) and the maximal rate *k*_0_ is achieved, whereas the promoter is mostly closed when DNA is strongly relaxed.

The main ingredient of the model, the opening energy ΔG_*op*_, is computed from an established coarse-grained unidimensional description of DNA twist-dependent thermodynamics [31, 32], where the total SC level is assumed to contribute to DNA opening by RNAP (neglecting any effect of its partitioning into twist/writhe and constrained/unconstrained contributions in the thermodynamic equilibrium of open-complex formation). Under this approximation, it is possible to predict the quantitative regulatory effect of SC variations from their experimentally available average value (e.g., using chloroquine-agarose gels), and its validity is justified *a posteriori* by the good agreement with *in vitro* and *in vivo* expression data (Figs. 1 and 2). Note that, at the genomic scale, the SC level locally available to RNAP for the opening of a given promoter may deviate from the genome-averaged SC level because of many complicating factors beyond the simple model considered here (3D conformation of the promoter, binding of regulatory proteins and NAPs, structural transitions occurring at nearby sites etc; see Discussion). But, because of the monotonous nature of the activation curves (Figs. 1 B and E), all main results are robust when the SC levels are globally shifted by up to *±*0.01.

For each promoter, the denaturation energy is computed with TwistDNA [32] using the 14-bp sequence starting from (and including) the −10 hexamer, corresponding to the extent of the transcription bubble (flanked by 100-bp long G-tracts to avoid boundary effects in the computation). The only adjustable parameter of TwistDNA is an effective salt concentration, which is calibrated on the data of Fig. 1 [20], yielding values of mM and 3 mM for *in vitro* and *in vivo* transcription, respectively, the latter value being kept constant for all subsequent *in vivo* calculations. These low values are likely due to the strongly simplified d escription o f the solvent (continuous distribution of monovalent ions) and DNA (unidimensional molecule) used in that software, and should thus be considered as effective parameters for the computation rather than quantitative concentrations.

### Superhelical densities

*In vivo* SC levels used in the computations of Fig. 1 were taken from [20, 16] (*E. coli* strains with norfloxacin), and [34] (topoisomerase mutants of *E. coli*). In the data of [19] (Fig. 1F), the activation levels are slightly higher than observed with *tyrT* and computed by the model, possibly due to competing structural transitions in the promoter sequence and/or tridimensional mechanical effects beyond our modelling.

Relaxation fold-changes measured in microplates with *pheP*, *pelE* and *pelD*-derived promoters were reproduced (Fig. 2) with a relaxation magnitude Δ*σ* = 0.001, starting from a level *σ* = −0.032 in LB rich medium, and *σ* = −0.023 for M63+G minimal medium. This low magnitude may be partly due to the slow growth conditions in microplates, but mostly to a buffering effect of the reporter system (luciferase lifetime of several to tens of minutes) and should thus be considered as an effective value used in the modelling, as also suggested by the low repressive effect of novobiocin compared to batch cultures [35].

For the computation of the genome-wide contribution to the relaxation-response (Discussion), transcription rates from all promoters are normalised by their sum in each condition before computing fold-changes (consistent with transcriptomic analysis protocols), resulting in the activation of a fraction of promoters (since the G/C-rich promoters represent a weaker proportion of total transcripts after the relaxation, A/T-rich promoters appear as activated, see Supplementary Fig. S6). Levels of SC variations associated to all investigated conditions were reviewed from the literature (Supplementary Tab. S2), exhibiting magnitudes in the range 0.01-0.015, with differences due to protocols in stress/shock conditions and chloroquine-agarose gel assays. Consequently, all model predictions were computed with an initial SC level *σ* = −0.045 (yielding the best overall agreement with observations) and a variation of Δ*σ* = ±0.015 (depending on the sign of the experimental response). The model predictions change only marginally when these figures are changed by less than 0.01.

## Acknowledgements

We thank Joanna Bonci and Nicolas Paulhan for experimental contributions, the whole CRP team for helpful discussions, Georgi Muskhelishvili and Ivan Junier for their critical reading of the manuscript, and a referee for sound criticisms that helped improving the manuscript.

