## Supplementary Information for "The discriminator sequence is a primary determinant in the supercoiling response of bacterial promoters"

May 18, 2021

**This PDF file includes:**

- Supplementary text
- Figures S1 to S6
- Tables S1 to S2
- SI references

### Supplementary text

#### Insertions of *pelD-luc* and *pelE-luc* transcriptional fusions in *D. dadantii* chromosome

All Plasmids and primers used for these constructions are described in Supplementary Tab. S1. *D. dadantii* 3937-derivative strains A5740 and A5720 with *pelD-luc* and *pelE-luc* transcriptional fusions, respectively, inserted in the *pelA-pelE* intergenic region of the chromosome, were obtained as follow. First, two PCR products of 500 bp, corresponding to the *pelA* and *pelE* genes flanking the site of insertion, were obtained using primer pairs *pelAF1/pelAR1* and *pelEF1/pelER1*. Primers *pelAR1* and *pelEF1* included a unique restriction site for BglII and were designed to have a 20-bp short overlapping complementary sequence. The two resulting PCR products were fused by overlapping PCR using primers *pelAF1* and *pelER1*. The resulting *pelA-BglII-pelE* PCR product was cloned into pGEMT plasmid (pGEMT-*pelA*-BglII-*pelE*). Then, PCR fragments containing either *pelD-luc-CmR* or *pelE-luc-CmR* were obtained from plasmids pUCTer-*pelD-luc* and pUCTer-*pelE-luc*, respectively, by using primer pairs C18 and 155 with a BglII restriction site at their 5' extremities. Finally, the pGEMT-*pelA*-BglII-*pelE* plasmid and the PCR *pelD-luc-CmR* or *pelE-luc-CmR* fragments were digested with BglII and ligated. After transformation in *E. coli*, plasmids containing the expected insertions were selected and electroporated into *D. dadantii* strain 3937 using a standard electroporation procedure. The two insertions were introduced into *D. dadantii* chromosome by marker exchange recombination between chromosomal and plasmid-borne alleles. The recombinants were selected after successive cultures in low phosphate medium in the presence of chloramphenicol, conditions in which pGEMT derivatives are very unstable [1]. The recombination was finally validated by PCR. *D. dadantii* strain 3937 cells were also transformed with plasmids carrying *pelE* and *pelD* native promoters (pUCTer-*pelE-luc* and pUCTer-*pelD-luc*, respectively, Supplementary Tab. S1) for further comparison with chromosomal fusions.

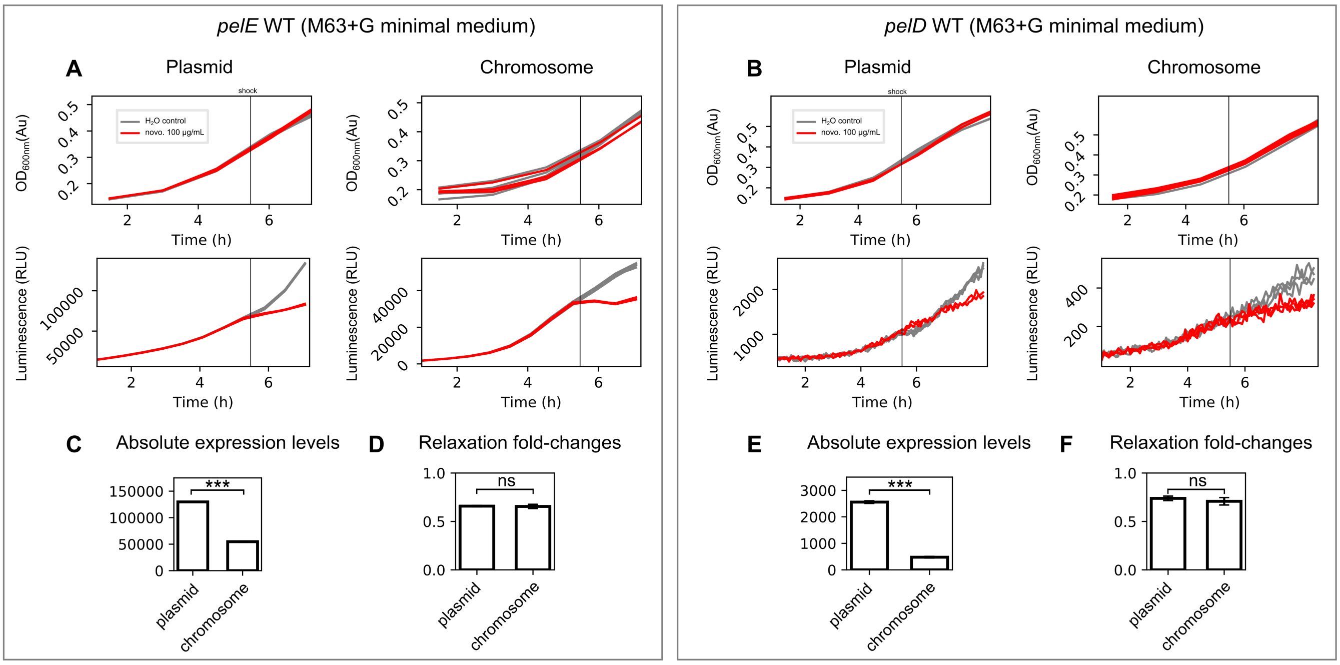


**Supplementary Figure S1**: Bacterial growth and promoter expression monitored in a microplate reader, for *pelE-luc* and *pelD-luc* transcriptional fusions either inserted into *D. dadantii* chromosome or incorporated into pUCTer-*luc* plasmids and transformed into *D. dadantii* cells. (A) Bacterial growth measured by $OD_{600nm}$ and *pelE* expression level measured by luminescence in minimal medium. A novobiocin shock was applied in mid-exponential phase ($100\mu g/mL$ concentration). (B) Same for *pelD*. The patterns of bacterial growth, promoter expression and response to novobiocin are consistent between the two systems (chromosomal fusions vs plasmid-borne fusions), for both *pelE* and *pelD*. (C) Absolute expression levels for *pelE* promoter in absence of shock: as expected, the multicopy plasmid-borne fusion is expressed at a higher level than the chromosome-borne fusion. (D) Comparison of relaxation fold-changes for *pelE* promoter, demonstrating a quantitatively identical effect of novobiocin in the two constructions, in spite of the difference of absolute expression. This observation confirms that the fold-changes observed on the plasmid-borne constructions are indeed due to a regulatory effect of DNA relaxation (identical in the chromosome and in the plasmid) rather than a plasmid-specific effect (copy number variation, etc). The relaxation activity of novobiocin was previously demonstrated in *D. dadantii* cells on the plasmids employed [2]. (E) and (F) same for *pelD* promoter. (C)-(F) were computed from the final luminescence time-points in (A) and (B).

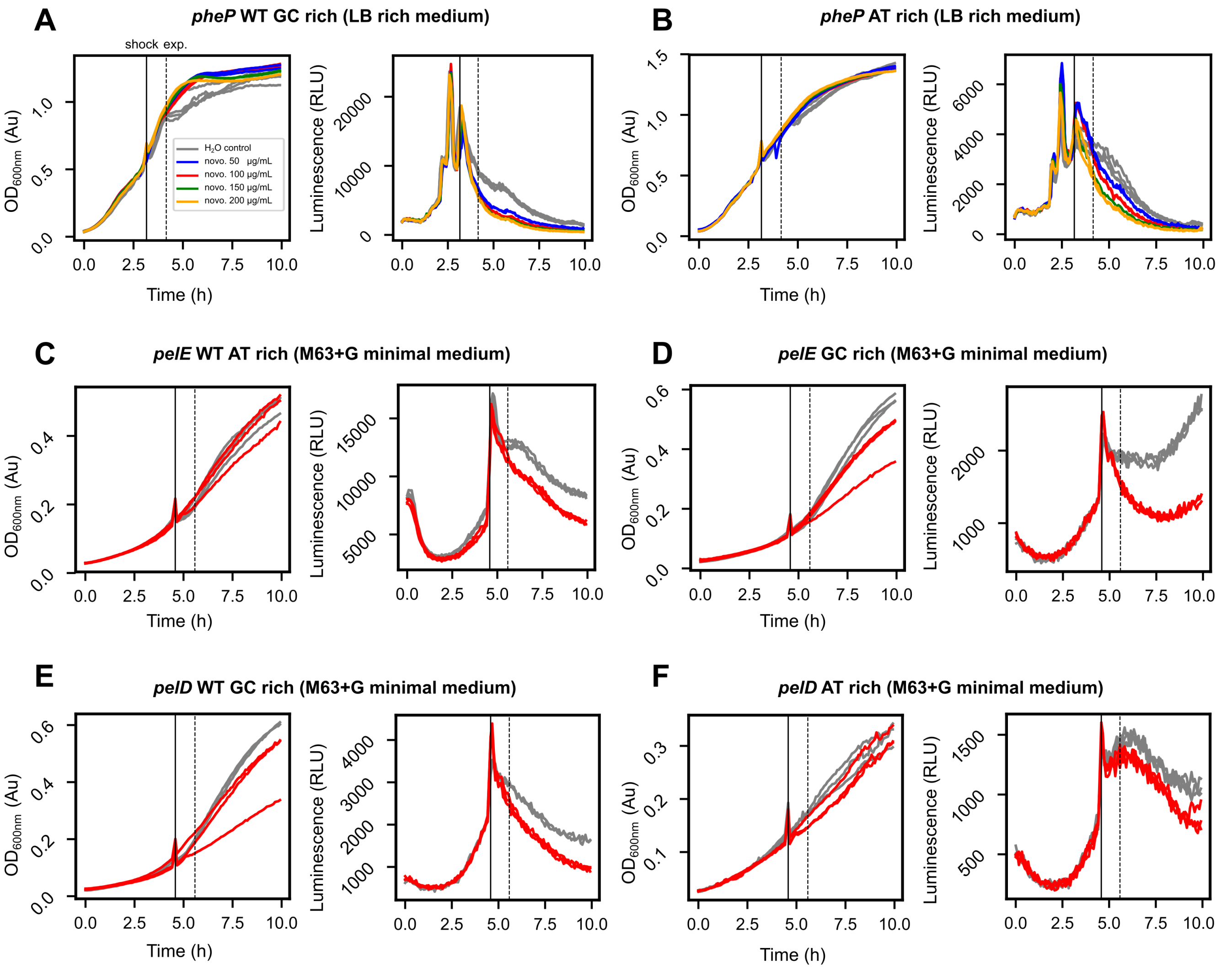


**Supplementary Figure S2**: Bacterial growth and mutant promoter expression monitored in *E. coli* in a microplate reader (raw data). For all promoters, the left panel shows bacterial growth measured by $OD_{600nm}$ and the right panel shows the expression level measured luminescence. (A) *E. coli* bacteria carrying plasmids with *pheP* WT promoter in rich medium. (B) *pheP* AT-rich promoter. (C) *pelE* WT promoter in minimal medium with only 100 µg/mL novobiocin concentration tested. (D) *pelE* GC-rich promoter. (E) *pelD* WT promoter. (F) *pelD* AT-rich promoter. The $OD_{600nm}$ discrepancies at shock time or after several hours in some cases (D, E, F) are optical artefacts due to the opening of the recorder and/or the formation of sediments disrupting the measurements (the luminescence does not vary correspondingly among replicates).

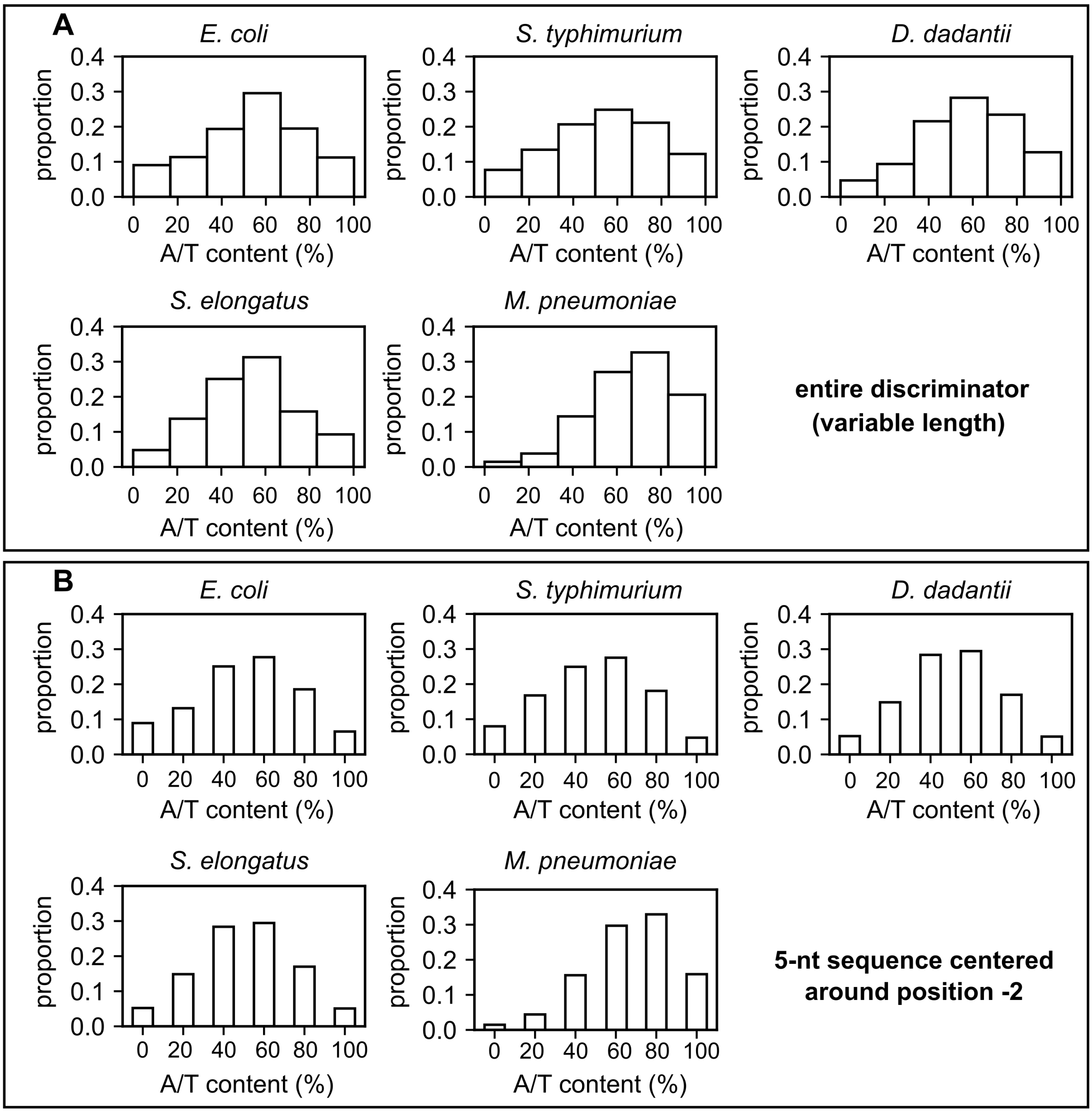


**Supplementary Figure S3**: Variability of promoter discriminator A/T content in the investigated organisms. Proportion of promoters depending on their A/T content in the discriminator defined as (A) the sequence of variable length between -10 element and +1 positions or (B) the 5-nt sequence centred around position -2 used throughout the manuscript. For *E. coli*, *S. typhimurium*, *D. dadantii* and *M. pneumoniae*, only $\sigma 70$ promoters were considered, whereas only $\sigma A$ promoters were considered for *S. elongatus*. All promoters were aligned at their -10 site, except for *S. elongatus* and *M. pneumoniae* for which promoters were aligned at their annotated TSS, and the signal shifted of 5 nt (see Materials and Methods). The distributions are similar with the two methods.

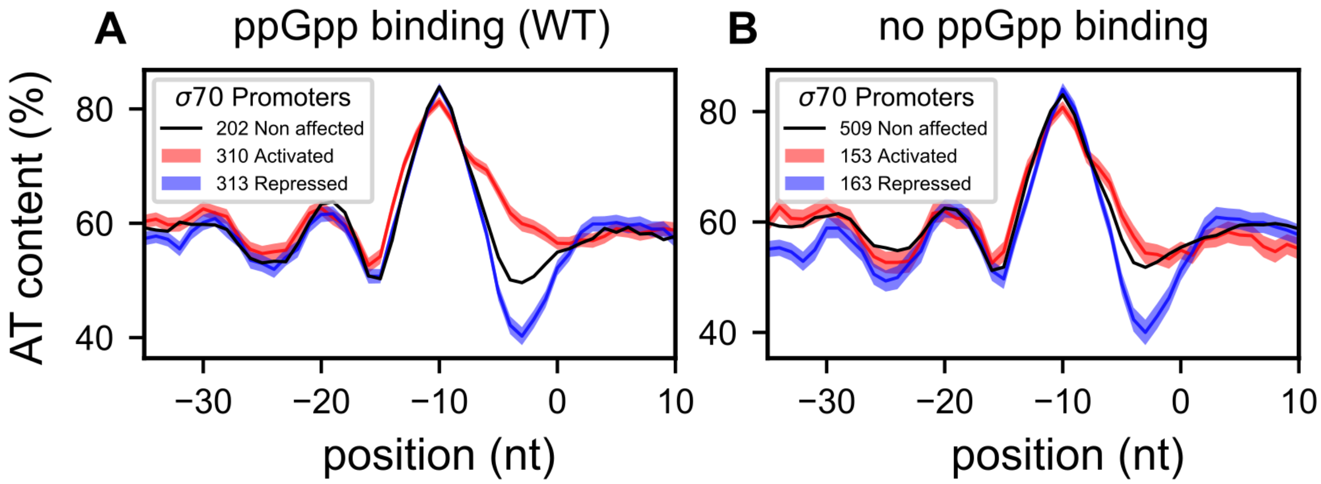


**Supplementary Figure S4**: Simultaneous regulation by SC and ppGpp at the discriminator. (A) Average A/T% profiles of *E. coli* $\sigma$70 promoters along 5-bp centred windows, depending on their response to ppGpp induction. Data from [3], strain 1-2-pALS13 after 5 minutes (310 activated and 313 repressed promoters). The A/T% profiles of the promoter groups are very similar (overlapping 67% confidence intervals, visible as coloured shaded areas) except in the discriminator region (between -10 and +1 positions). (B) Same in the 1-2-pALS13 strain after 10 minutes harbouring a mutant RNA Polymerase unable to bind ppGpp. ppGpp still induces a differential expression of around half as many genes (153 activated and 160 repressed promoters), and the difference in A/T% is still strongly present, although weaker than in A. We suggest ppGpp-induced SC relaxation as a plausible mechanism for this regulation.

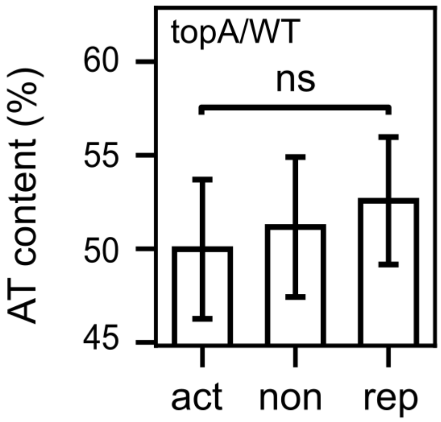


**Supplementary Figure S5**: A/T% in the discriminator of promoters differentially expressed in a *topA* mutant strain (vs WT), associated to an inheritable increase in SC. G/C-rich discriminators tend to be more activated in the *topA* strain compared to the WT strain as expected from our modelling, although the results are not significant (P=0.18). This mutation alone can therefore not explain the stronger signal observed in the evolution experiment (Fig. 6).

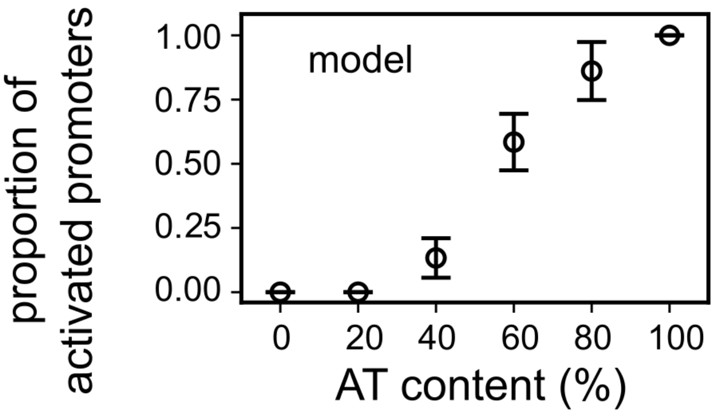


**Supplementary Figure S6**: Model prediction of the response of *S. enterica* promoters to DNA relaxation. After applying the model of Eq. 1 to all promoters, a normalisation step mimics the protocol of the transcriptomic experiment. The proportion of promoters predicted as activated is then computed depending on their A/T% in a 5-nt window around position -2 (same as Fig. 3B and D).

| Promoter | Sequence |
|---|---|
| >*pheP_WT_GCrich* | CTCGAGTCAGAGGTGATGAGCCGGATTGCCGCGCCGATGATTGGCGGCATGATCACCGCACCTTTGCTGTCGCTGTTTATT<br>ATCCCGGCGGCGTATAAGCTGATGTGGCTGCACCGACATCGGGTACGGAAATAAAAGCAGGATACCCCGTTTAACCGTGTG<br>GATTGTGTCTTGCGACGATGGGCACTAAATGTTAAAAG**GTGCCC**CTCAACAAAAAAGACACACAGGGGAAAGGCGGATCC |
| >*pheP_hybrid* | CTCGAGTCAGAGGTGATGAGCCGGATTGCCGCGCCGATGATTGGCGGCATGATCACCGCACCTTTGCTGTCGCTGTTTATT<br>ATCCCGGCGGCGTATAAGCTGATGTGGCTGCACCGACATCGGGTACGGAAATAAAAGCAGGATACCCCGTTTAACCGTGTG<br>GATTGTGTCTTGCGACGATGGGCACTAAATGTTAAAAG**GTTACC**CTCAACAAAAAAGACACACAGGGGAAAGGCGGATCC |
| >*pheP_ATrich* | CTCGAGTCAGAGGTGATGAGCCGGATTGCCGCGCCGATGATTGGCGGCATGATCACCGCACCTTTGCTGTCGCTGTTTATT<br>ATCCCGGCGGCGTATAAGCTGATGTGGCTGCACCGACATCGGGTACGGAAATAAAAGCAGGATACCCCGTTTAACCGTGTG<br>GATTGTGTCTTGCGACGATGGGCACTAAATGTTAAAAG**GTTAAT**CTCAACAAAAAAGACACACAGGGGAAAGGCGGATCC |
| >*pelE_WT_ATrich* | CTCGAGTCGAAATTAAAAATATAAATAATTGATTTAAATCATAAAAAATAAAAAACCAAGTAACACAAAGTTACAAATACA<br>GTCAATAGTTTATTTTTTATTAATAAAAACCATTGTCATCATCGTGACAAAGTTCACAAAATAGACACTCAAACCGCATAA<br>ACATTCGCCAAAGCAAAAGATGAAATGGTATTTC**GTTTT**TAGACACACATGTAACAAATGGACACCATTTGGATCGCTCAC<br>TGAGCACACAAGGAAAATTGCCATGAACAACTCACGTATGTCTTCCGTTTCAACACAGAAAACAACAGGACGTTCTGCCTT<br>GGGAACCAAAGGGATCC |
| >*pelE_GCrich* | CTCGAGTCGAAATTAAAAATATAAATAATTGATTTAAATCATAAAAAATAAAAAACCAAGTAACACAAAGTTACAAATACA<br>GTCAATAGTTTATTTTTTATTAATAAAAACCATTGTCATCATCGTGACAAAGTTCACAAAATAGACACTCAAACCGCATAA<br>ACATTCGCCAAAGCAAAAGATGAAATGGTATTTC**GCGCC**TAGACACACATGTAACAAATGGACACCATTTGGATCGCTCAC<br>TGAGCACACAAGGAAAATTGCCATGAACAACTCACGTATGTCTTCCGTTTCAACACAGAAAACAACAGGACGTTCTGCCTT<br>GGGAACCAAAGGGATCC |
| >*pelD_WT_GCrich* | CTCGAGAACTGTTTGGGTTATTTCAGATAAAAACGCTTATACATATAGCTGAATTTAAAAGAAAAATTAATTCAACATTCA<br>TAACTAAAAGTTACCGGTCACGATCACACTTTAGATAAAATTAATTAGCCTCATAAAAAAAACGAGATTTTGATCACAAAAT<br>AAACAATCGAAAACGCTTAAAAAT**CCGCC**TTGCCAAAGGACAAAATGGCGTTTCATTTTTTTCACAAACACTTTTCAGTC<br>AACAAAATTGGATTAGCGCAGATAGCGCAAGGAACCAGTCTATGAACAACACACGAGTGTCTTCCGTAGGTACCAAGGATCC |
| >*pelD_ATrich* | CTCGAGAACTGTTTGGGTTATTTCAGATAAAAACGCTTATACATATAGCTGAATTTAAAAGAAAAATTAATTCAACATTCA<br>TAACTAAAAGTTACCGGTCACGATCACACTTTAGATAAAATTAATTAGCCTCATAAAAAAAACGAGATTTTGATCACAAAAT<br>AAACAATCGAAAACGCTTAAAAAT**CTTTT**TTGCCAAAGGACAAAATGGCGTTTCATTTTTTTCACAAACACTTTTCAGTC<br>AACAAAATTGGATTAGCGCAGATAGCGCAAGGAACCAGTCTATGAACAACACACGAGTGTCTTCCGTAGGTACCAAGGATCC |
| | XhoI restriction site, -35 element, -10 element, **discriminator**, TSS, BglII restriction site |

| Plasmid | Description | Origin |
|---|---|---|
| pGEMT | High-copy-number vector containing a multiple cloning site within the alpha-peptide coding region of the enzyme beta-galactosidase. | Promega |
| pGEMT-*pelA*-BglII-*pelE* | pGEMT derivative containing both the 500-bp region with *pelA-BglII* and the 500-bp region with *pelE*. | This work |
| pUCTer-*luc* | High-copy-number vector (pUC18 derivative) containing a multiple cloning site upstream of the *luc* reporter gene, followed by a *rrnB* terminator and a *cat* gene conferring chloramphenicol resistance. | Laboratory collection |
| pUCTer-*pelD-luc* | pUCTer-*luc* derivative containing the *D. dadantii pelD* WT promoter sequence above cloned upstream of the *luc* reporter gene. | This work |
| pUCTer-*pelE-luc* | pUCTer-*luc* derivative containing the *D. dadantii pelE* WT promoter sequence above cloned upstream of the *luc* reporter gene. | This work |
| **Primer name** | **Sequence** | |
| *pelA* F1 | CTCAGGATAAAGGTAAGCTGC | |
| *pelA* R1 | AGATCTGATGACGGTGTGGGCTAGACG | |
| *pelE* F1 | CGTCTAGCCCACACCGTCATCAGATCTCGCCCGACTCGTCCCCTTTC | |
| *pelE* R1 | GGAAGCGACTGAGACCATCATG | |
| pUCTer C18 | GGGAGATCTACGACGTTGTAAAACGACGG | |
| pUCTer 155 | GGGAGATCTAAAAGGCCATCCGTCAGGATGGCCTTCTCCGGGTCGAATTTGCTTTCG | |
| | rrnBT2 terminator sequence, BglII restriction site | |

**Supplementary Table S1**: List of synthetic promoter sequences with mutated discriminators. List of plasmids and primers used to construct *D. dadantii* 3937-derivative strains with *pelD-luc* and *pelE-luc* transcriptional fusions in the *pelA-pelE* intergenic region of the chromosome.

| Species | Condition | Condition reference | SC variation | SC variation reference | Transcription start sites reference | A/T% difference p-value (act vs rep) | Model sensitivity gain (p-value) |
|---|---|---|---|---|---|---|---|
| *Salmonella typhimurium* | novobiocin | [4] | - | no measurement | [5] | $< 10^{-5}$ | 13.8% ($< 10^{-6}$) |
| *Dickeya dadantii* | novobiocin | [6] | - | [2] | [7] | $< 0.001$ | 9.8% ($< 0.001$) |
| *Escherichia coli* | norfloxacin | [8] | - | [8] | [9] | 0.010 | 5.1% (0.039) |
| *Synechococcus elongatus* | correlation* | [10] | - | [10] | [11] | 0.004 | 4.7% (0.099) |
| *Mycoplasma pneumoniae* | novobiocin | [12] | - | no measurement | [12] | 0.029 | 7.8% (0.020) |
| *Escherichia coli* | heat shock | [13] | - | [14] | [9] | $< 10^{-5}$ | 7.5% ($< 10^{-4}$) |
| | cold shock | [15] | + | [16] | [17] | $< 10^{-4}$ | 4.1% (0.038) |
| | oxidative shock | [15] | - | [18] | [9] | 0.017 | 9.0% (0.013) |
| *Dickeya dadantii* | oxidative shock | [19] | - | [2] | [7] | $< 10^{-4}$ | 7.6% ($< 0.001$) |
| *Escherichia coli* | experimental evolution (2K mutant) | [6] | + | [20] | [9] | 0.005 | 3.7% ((0.011) |
| | experimental evolution (20K mutant) | [6] | + | [20] | [9] | 0.011 | 1.8% (0.18) |

**Supplementary Table S2**: Compilation of investigated species, conditions and results. The SC change refers to the direction of SC variation induced by the condition (-: DNA relaxation, +: increase in negative SC). The correlation* condition from *S. elongatus* corresponds to the phasing of gene expression in the SC circadian oscillation and provides an indirect proxy of gene response to SC relaxation [10]. For the stress conditions, the protocols used in the SC assays usually differ from those used in the transcriptomic experiments, and thus only give a semi-quantitative estimate of the levels involved in the observed regulatory response. A/T% contents are compared with Student tests, in 5-nt windows centred at position -2. The sensitivity gain is computed as the increase in the proportion of accurately predicted genes (among significantly activated or repressed genes) compared to a null (random) model. Additional detail in Materials and Methods.
